# Chromosome-scale haplotype-phased genome assemblies of the male and female lines of wild asparagus (*Asparagus kiusianus*), a dioecious plant species

**DOI:** 10.1101/2021.12.15.472771

**Authors:** Kenta Shirasawa, Saki Ueta, Kyoko Murakami, Mostafa Abdelrahman, Akira Kanno, Sachiko Isobe

## Abstract

*Asparagus kiusianus* is a disease-resistant dioecious plant species and a wild relative of garden asparagus (*A. officinalis*). To enhance *A. kiusianus* genomic resources, advance plant science, and facilitate asparagus breeding, we determined the genome sequences of the male and female lines of *A. kiusianus*. Genome sequence reads obtained with a linked-read technology were assembled into four haplotype-phased contig sequences (~1.6 Gb each) for the male and female lines. The contig sequences were aligned onto the chromosome sequences of garden asparagus to construct pseudomolecule sequences. Approximately 55,000 potential protein-encoding genes were predicted in each genome assembly, and ~70% of the genome sequence was annotated as repetitive. Comparative analysis of the genomes of the two species revealed structural and sequence variants between the two species as well as between the male and female lines of each species. Genes with high sequence similarity with the male-specific sex determinant gene in *A. officinalis, MSE1/AoMYB35/AspTDF1*, were presented in the genomes of the male line but absent from the female genome assemblies. Overall, the genome sequence assemblies, gene sequences, and structural and sequence variants determined in this study will reveal the genetic mechanisms underlying sexual differentiation in plants, and will accelerate disease-resistance breeding in garden asparagus.

## Introduction

*Asparagus kiusianus* is a wild relative of garden asparagus (*Asparagus officinalis*). While garden asparagus is a cultivated species belonging to the Asparagaceae family and is consumed as a vegetable crop around the world, *A. kiusianus* is native to the coastal regions of Japan^1^. Therefore, *A. kiusianus* might exhibit tolerances and/or resilience to abiotic and biotic stresses. Although *A. kiusianus* has been identified as a potential donor of stem-blight disease resistance in asparagus breeding programs^2^, neither the genetic mode nor the genetic loci of resistance have been elucidated to date.

*A. officinalis* is a dioecious species and is widely recognized as a model for sex determination in plants. Recent studies indicate that the male-specific *MYB*-like gene, *MSE1/AoMYB35/AspTDF1*, located at the masculinization-promoting *M* locus of the Y-specific region in asparagus, functions in sex determination in asparagus^3–5^. Since *A. kiusianus* is also a dioecious plant species, like garden asparagus, it is possible that both species share the same system of sex determination. Therefore, comparative genome sequence and structure analyses between the two species could provide insights into the molecular mechanisms underlying sex determination in Asparagaceae and the evolutionary processes involved therein.

Advances in sequencing technologies have enabled the whole-genome sequencing of various plant species, thus providing fundamental information required for understanding the plant biology and accelerating breeding programs. Nevertheless, while the genome sequence data of garden asparagus^3^ and transcriptome data of *A. kiusianus*^6,7^ have been made publicly available, no whole-genome sequence data have been released for *A. kiusianus* to date. Owing to the dioecious nature of *A. kiusianus*, which leads to allogamy, its genome is predicted to be highly heterozygous. Therefore, haplotype-phased genome sequence data would be useful for dissecting the allelic sequence and structural variations in *A. kiusianus*. In this study, we employed a linked-read technology (10X Genomics, Pleasanton, CA, USA) to construct haplotype-based genome sequence assemblies of the male and female lines of *A. kiusianus*. The genome sequence assemblies were then used for gene prediction and sequence and structural variant discovery. Overall, the genome sequence information of *A. kiusianus* obtained in this study could accelerate studies on plant sex determination and facilitate asparagus breeding programs.

## Materials and methods

### Plant materials

Male (K1) and female (K2) lines of *A. kiusianus* cultivated at Kagawa Prefectural Agricultural Experiment Station (Kagawa, Japan) were used in this study. Genomic DNA was extracted from the stems of young seedlings using the modified cetyltrimethylammonium bromide (CTAB) method^8^.

### Genome sequencing and assembly

Genomic DNA libraries of male and female lines were prepared using the Chromium Genome Library Kit v2 (10X Genomics), and sequenced on NovaSeq 6000 (Illumina, San Diego, CA, USA) in paired-end, 150 bp mode. The sequence reads were assembled with Supernova (10X Genomics) to construct the contig sequences, scaffold the contigs, and resolve haplotype phases. DNA library preparation, sequencing, and assembly were conducted by Takara Bio (Shiga, Japan) as an outsourcing service.

The genome sizes of male and female lines were estimated based on short reads using Jellyfish. To construct pseudomolecule sequences at the chromosome level, the assembled contigs were aligned against the sequence of 10 *A. officinalis* chromosomes (reference) using RaGoo.

The software tools used for data analyses are listed in Supplementary Table S1.

### Repetitive sequence analysis and gene prediction

Repetitive sequences in the assemblies were identified with RepeatMasker, using repeat sequences registered in Repbase and a *de novo* repeat library built with RepeatModeler.

RNA-Seq reads of *A. kiusianus* and *A. officinalisi* were obtained from a public DNA database (GenBank Sequence Read Archive accession number: SRA1003110)^6,7^. The RNA-Seq reads, from which adapter sequences were trimmed with fastx_clipper in the FASTX-Toolkit, were aligned against the assembled sequences with HISAT2. Gene prediction was performed with BREAKER2 using the positional information of the repeats, RNAs, and peptide sequences of the predicted genes of *A. officinalis* (V1.1)^3^ released in Phytozome.

### *Comparative genome structure analysis of* A. kiusianus *and* A. officinalis

Chromosome-level genome sequence assemblies of *A. kiusianus* (this study) and *A. officinalis* (V1.1)^3^ were compared with Minimap2, and the resultant Pairwise mApping Format (PAF) files were visualized with pafr.

## Results and data description

### Haplotype-phased genome assembly

Short-read sequences of the male (143.7 Gb) and female (140.0 Gb) lines of *A. kiusianus* were obtained in this study, and their genome sizes were estimated at 1,563.8 and 1,729.4 kb, respectively (Figure 1).

**Figure 1.**
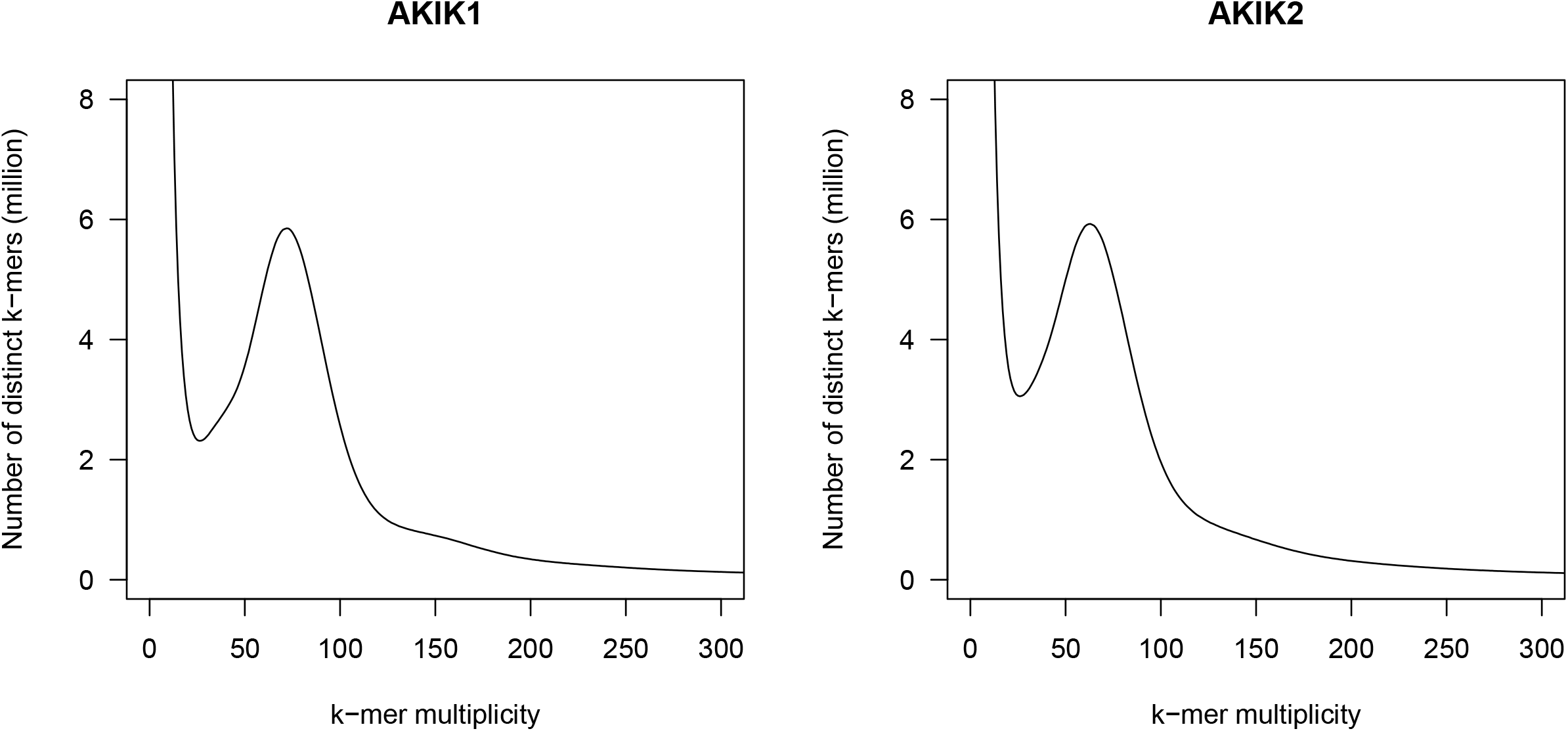
Estimation of the genome sizes of the male and female lines of *Asparagus kiusianus*, based on *k*-mer analysis (*k* = 17) with the given multiplicity values.

The short-read sequences of the male line were assembled into raw contigs (total length = 3,724.6 kb, N50 = 7.5 kb), which included gaps and all homologous sequences of the diploid genome (Supplementary Table S2). Then, the homologous sequences were flattened, and the gaps were filled by joining the sequence to its flanking sequence, thus producing megabubble sequences (total length = 1,811.6 kb, N50 = 170.6 kb) (Supplementary Table S2). Finally, two haplotype-phased genome assemblies (each containing 111,443 sequences) were generated from the megabubble sequences (Table 1). Haplotype 1 spanned 1,618.9 Mb in total with an N50 value of 155.5 kb, while haplotype 2 spanned 1,618.5 Mb in total with an N50 length of 155.3 kb. Complete Benchmarking Single-Copy Orthologs (BUSCO) scores were 88.4% and 88.6% for haplotypes 1 and 2, respectively (Table 1). The male genome assemblies for haplotype 1 and 2 were designated as AKIK1p1 and AKIK1p2, respectively.

**Table 1.** Statistics of the genome assemblies of the male and female lines of *Asparagus kiusianus*.

|  | AKIK1p1_r1.0 | AKIK1p2_r1.0 | AKIK2p1_r1.0 | AKIK2p2_r1.0 |
| --- | --- | --- | --- | --- |
| Total contig size (bp) | 1,618,866,837 | 1,618,519,636 | 1,567,596,515 | 1,567,276,205 |
| Number of sequences | 111,443 | 111,443 | 125,241 | 125,241 |
| Sequence N50 length (bp) | 155,490 | 155,269 | 59,656 | 59,610 |
| Longest sequence size (bp) | 2,375,321 | 2,375,321 | 1,231,068 | 1,222,162 |
| Gap size (bp) | 168,294,890 | 168,291,350 | 132,162,460 | 132,161,290 |
| Single-copy complete BUSCOs | 82.4% | 82.6% | 83.5% | 83.4% |
| Duplicated complete BUSCOs | 6.0% | 6.0% | 5.2% | 5.2% |
| Fragmented BUSCOs | 5.8% | 5.5% | 5.6% | 5.6% |
| Missing BUSCOs | 5.8% | 5.9% | 5.7% | 5.8% |

Similarly, short-read sequences of the female line were assembled into raw contigs (total length = 3,780.7 kb, N50 = 7.0 kb) and megabubble sequences (total length = 1,703.6 Gb, N50 = 76.2 kb) (Supplementary Table S2). The resultant haplotype-phased assemblies (each containing 125,241 sequences) spanned 1,567.6 Mb in total with an N50 value of 59.7 kb for “haplotype 1”, and 125,241 sequences of 1,567.3 Mb in total with an N50 value of 59.6 kb for “haplotype 2” (Table 1). Complete BUSCO scores were 88.7% and 88.6% for haplotypes 1 and 2, respectively (Table 1). The female genome assemblies for haplotype 1 and 2 were designated as AKIK2p1 and AKIK2p2, respectively.

The four sets of genome sequence assemblies of *A. kiusianus* (haplotypes 1 and 2 of male and female lines) were aligned against the chromosome-scale genome assembly of *A. officinalis*. A total of 96,224 sequences (1,535.9 kb) for haplotype 1 and 96,224 sequences for haplotype 2 (1,535.3 kb) in the male line, and 107,875 sequences (1,491.7 kb) for haplotype 1 and 107,864 sequences (1,489.7 kb) for haplotype 2 in the female line, could be aligned to the 10 chromosome sequences of *A. officinalis* (Table 2).

**Table 2.** Statistics of the pseudomolecule sequences of *A. kiusianus*.

|  | AKIK1p1_r1.0 |  |  |  | AKIK1p2_r1.0 |  |  |  |
| --- | --- | --- | --- | --- | --- | --- | --- | --- |
|  | Total length (bp) | Gap (%) | Number of contigs | Number of genes | Total length (bp) | Gap (%) | Number of contigs | Number of genes |
| ch01 | 179,686,458 | 11.8 | 10,537 | 6,938 | 179,663,709 | 11.8 | 10,543 | 6,706 |
| ch02 | 94,379,980 | 12.5 | 6,056 | 3,631 | 94,286,185 | 12.5 | 6,066 | 3,427 |
| ch03 | 160,690,220 | 10.4 | 10,792 | 6,135 | 161,086,385 | 10.5 | 10,791 | 5,855 |
| ch04 | 222,888,549 | 11.1 | 13,146 | 7,547 | 222,850,935 | 11.1 | 13,139 | 7,251 |
| ch05 | 175,875,079 | 12.0 | 10,573 | 6,878 | 176,035,479 | 12.0 | 10,582 | 6,668 |
| ch06 | 112,254,750 | 9.4 | 7,553 | 3,696 | 112,182,592 | 9.4 | 7,553 | 3,504 |
| ch07 | 231,128,771 | 11.0 | 13,973 | 8,285 | 230,178,869 | 11.0 | 13,928 | 7,953 |
| ch08 | 209,954,455 | 10.5 | 13,763 | 7,167 | 209,720,314 | 10.5 | 13,796 | 6,882 |
| ch09 | 73,375,337 | 10.6 | 4,435 | 2,759 | 73,241,043 | 10.7 | 4,437 | 2,638 |
| ch10 | 85,316,880 | 10.8 | 5,396 | 3,353 | 85,678,944 | 10.9 | 5,389 | 3,229 |
| **Subtotal** | **1,545,550,479** | **11.0** | **96,224** | **56,389** | **1,544,924,455** | **11.0** | **96,224** | **54,113** |
| ch00 | 84,496,458 | 10.4 | 15,223 | 2,819 | 84,775,281 | 10.6 | 15,223 | 2,593 |
| **Total** | **1,630,046,937** | **11.0** | **111,447** | **59,208** | **1,629,699,736** | **11.0** | **111,447** | **56,706** |

|  | AKIK2p1_r1.0 |  |  |  | AKIK2p2_r1.0 |  |  |  |
| --- | --- | --- | --- | --- | --- | --- | --- | --- |
|  | Total length (bp) | Gap (%) | Number of contigs | Number of genes | Total length (bp) | Gap (%) | Number of contigs | Number of genes |
| ch01 | 167,748,676 | 9.5 | 11,500 | 6,772 | 167,821,022 | 9.5 | 11,527 | 6,835 |
| ch02 | 88,515,862 | 10.3 | 6,726 | 3,557 | 87,905,216 | 10.2 | 6,698 | 3,651 |
| ch03 | 159,691,030 | 9.0 | 12,193 | 5,956 | 159,408,608 | 9.0 | 12,181 | 6,030 |
| ch04 | 217,008,733 | 9.0 | 15,036 | 7,377 | 217,068,856 | 9.0 | 15,080 | 7,655 |
| ch05 | 172,460,497 | 9.9 | 11,974 | 6,762 | 171,835,019 | 9.9 | 11,943 | 6,970 |
| ch06 | 107,102,574 | 8.1 | 8,296 | 3,565 | 107,106,266 | 8.1 | 8,278 | 3,621 |
| ch07 | 222,804,915 | 9.2 | 15,815 | 7,998 | 221,855,507 | 9.2 | 15,811 | 8,158 |
| ch08 | 213,448,917 | 8.6 | 15,408 | 7,141 | 213,032,411 | 8.6 | 15,401 | 7,250 |
| ch09 | 71,157,753 | 9.4 | 5,055 | 2,517 | 71,672,067 | 9.5 | 5,054 | 2,573 |
| ch10 | 82,583,459 | 9.3 | 5,872 | 3,262 | 82,780,279 | 9.2 | 5,891 | 3,270 |
| **Subtotal** | **1,502,522,416** | **9.2** | **107,875** | **54,907** | **1,500,485,251** | **9.2** | **107,864** | **56,013** |
| ch00 | 77,633,199 | 8.8 | 17,366 | 2,616 | 79,350,054 | 8.9 | 17,377 | 2,681 |
| **Total** | **1,580,155,615** | **9.2** | **125,241** | **57,523** | **1,579,835,305** | **9.2** | **125,241** | **58,694** |

### Repetitive sequence analysis and gene prediction

Repeat sequences occupied 66.8% (AKIK1p1), 66.8% (AKIK1p2), 68.1% (AKIK2p1), and 68.1% (AKIK2p2). The most abundant repetitive sequences were long terminal repeats (LTRs) (45.4–46.3%), followed by unclassified repeats (14.9–13.7%) and DNA transposons (4.7–4.8%) (Table 3).

**Table 3.**
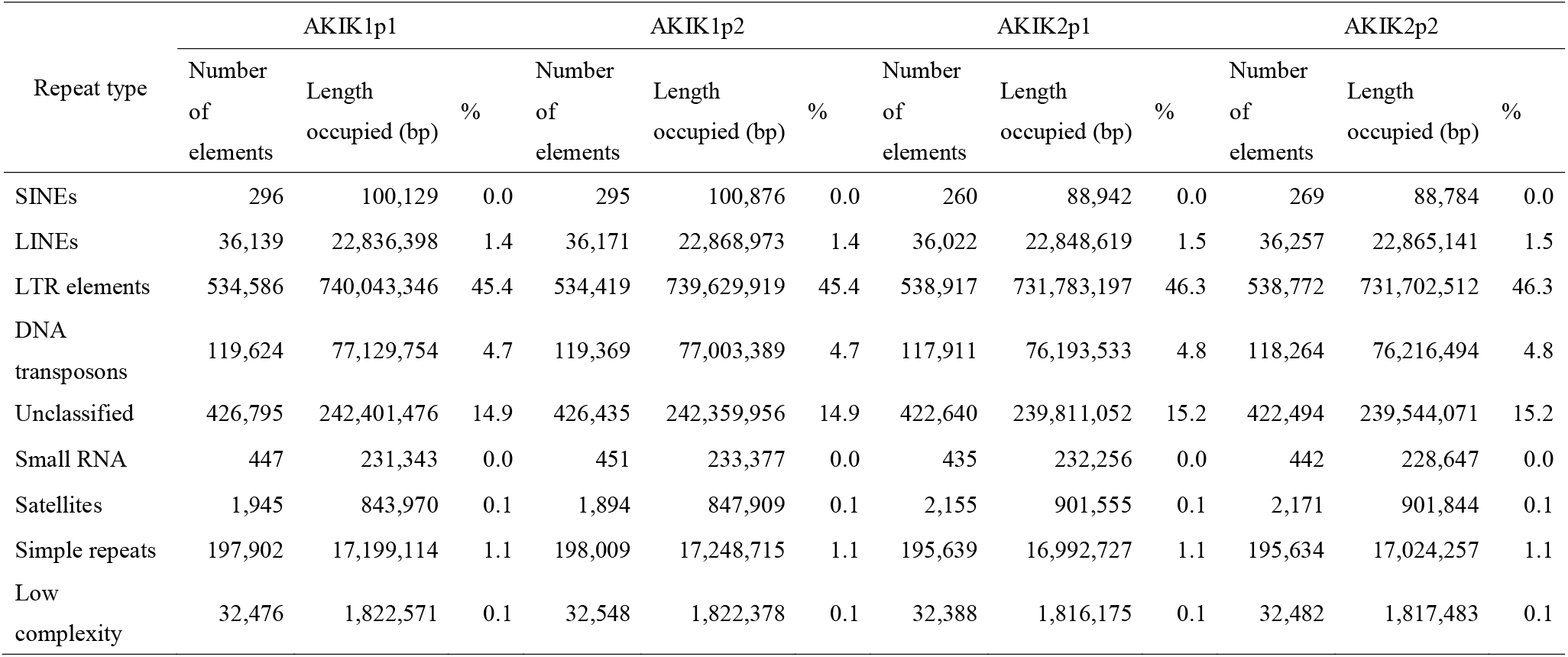
Repetitive sequences in the *A. kiusianus* genomes.

A total of 404.2 million RNA reads for 18 samples were mapped to the genome sequences. The mapping rates of *A. kiusianus* RNA-Seq reads were 93.7–94.1%, while those of *A. officinalis* reads were 82.4–82.6%. Based on the positions of RNA-Seq reads on the genome sequences, a total of 59,208, 56,706, 57,523, and 58,694 potential protein-coding genes were predicted in AKIK1p1, AKIK1p2, AKIK2p1, and AKIK2p2, respectively (Table 2), of which 365, 380, 472, and 505 genes contained premature termination codons in their internal sequences. Complete BUSCO scores ranged from 90.1% (AKIK2p1) to 91.4% (AKIK1p2).

Next, we compared the sequences of predicted genes with *MSE1/AoMYB35/AspTDF1*, which has been reported as the male-specific sex determinant gene in *A. officinalis*^3-5^. Two genes, K1p1ch01g28074 and K1p2ch01g47751, identified in the genomes of the male line exhibited high sequence similarity with the query; however, none of the genes in the female genome assemblies showed significant sequence similarity with the query.

### *Genome sequence and structural variations between* A. kiusianus *and* A. officinalis

Haplotype sequences within the male and female lines were compared. A total of 386,723 single nucleotide polymorphisms (SNPs; transition [Ts]/transversion [Tv] ratio = 3.0) and 46,007 insertions/deletions (indels) were identified between the two haplotypes of the male line (Table 4). On the other hand, 293,196 SNPs (Ts/Tv = 3.0) and 35,732 indels were identified between the two haplotype sequences of the female line (Table 4). The haplotype sequences of male and female lines were also compared, and 321,334 SNPs and 49,921 indels on average were identified across the four haplotype combinations (Table 4).

**Table 4.** Completeness of gene predictions, based on BUSCO assessment.

|  | Within a line |  | Between male and female lines |  |  |  |
| --- | --- | --- | --- | --- | --- | --- |
|  | AKIK1p1 | AKIK2p1 | AKIK1p1 | AKIK1p1 | AKIK1p2 | AKIK1p2 |
|  | vs | vs | vs | vs | vs | vs |
|  | AKIK1p2 | AKIK2p2 | AKIK2p1 | AKIK2p2 | AKIK2p1 | AKIK2p2 |
| Base substitutions | 386,723 | 293,196 | 324,361 | 327,460 | 314,704 | 318,810 |
| Ts/Tv ratio | 3.0 | 3.0 | 2.9 | 2.9 | 2.9 | 2.9 |
| 1bp indels | 23,833 | 17,283 | 20,344 | 20,379 | 19,692 | 19,970 |
| 2bp indels | 5,799 | 4,477 | 5,722 | 5,763 | 5,645 | 5,691 |
| [3,50) indels | 11,100 | 8,641 | 13,931 | 13,897 | 13,647 | 13,777 |
| [50,1000) indels | 1,871 | 1,472 | 7,141 | 7,127 | 7,096 | 7,096 |
| >=1000 indels | 3,404 | 3,859 | 3,199 | 3,229 | 3,160 | 3,179 |

While the chromosome structures were conserved within *A. kiusianus* lines and between *A. kiusianus* and *A. officinalis* (Figure 2), genomic rearrangements were observed at the local level. For instance, at the sex-related region including the male-specific gene *MSE1/AoMYB35/AspTDF1* of *A. officinalis*, sequence collinearity was disrupted by inversions and translocations between the male and female lines (Figure 3). Although sequence similarity was low between the male haplotypes of *A. kiusianus* and *A. officinalis*, sequence collinearity was moderately conserved (Figure 3).

**Figure 2.**
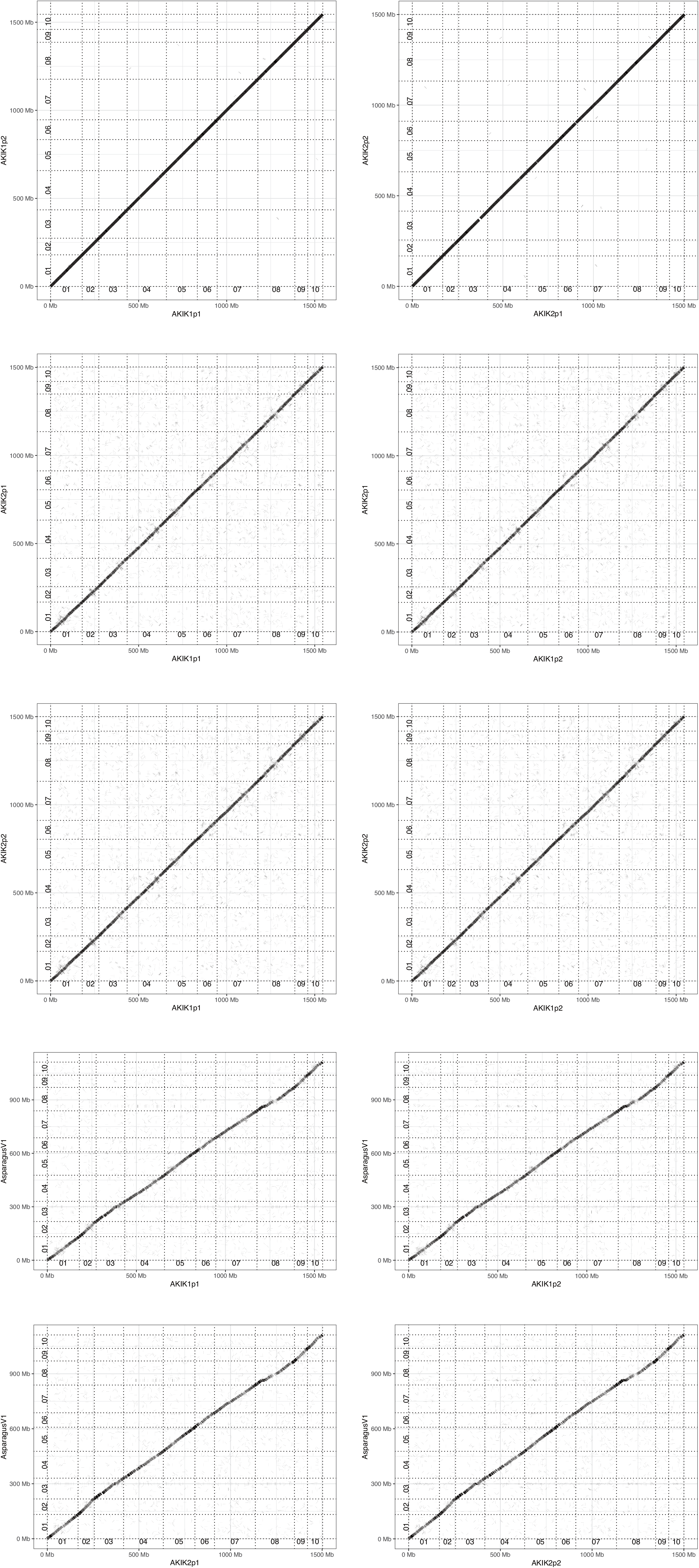
Comparative analysis of the genome sequence and structure of *Asparagus kiusianus* and *A. officinalis*. Dots represent similarities between the genome sequence and structure of the two species. Genomes of the male and female lines of *A. kiusianus* are indicated as AKIK1 and AKIK2, respectively. Haplotype phases 1 and 2 are indicated as p1 and p2, respectively. AsparagusV1 indicates *A. officinalis* genome.

**Figure 3.**
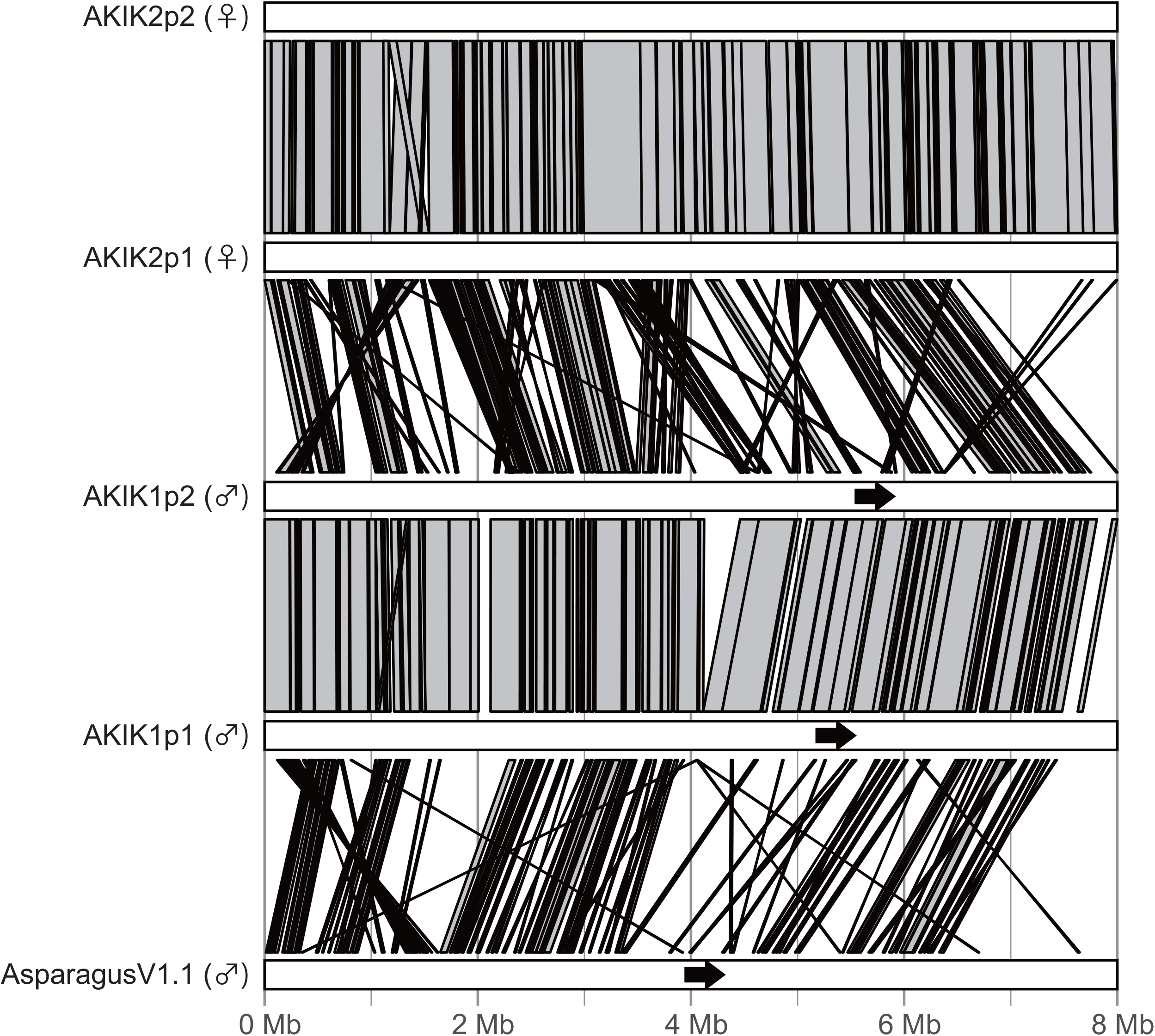
Comparison between the *M* locus in the Y-specific region of the male line and the corresponding region of the female line. Gray bars indicate similarities in the genome sequence and structure. Black arrows indicate the position and orientation of *MSE1/AoMYB35/AspTDF1*.

## Conclusion and future perspectives

We present the chromosome-level haplotype-phased genome assemblies of the male and female lines of *A. kiusianus*, a wild relative of garden asparagus. The genome size of *A. kiusianus* was estimated to be approximately 1.6 Gb (Figure 1), which was 300 Mb larger than that of garden asparagus (ca. 1.3 Gb)^3^. This estimation was reflected in the difference between the assembly sizes of *A. kiusianus* (1.6 Gb; Table 1) and garden asparagus (1.2 Gb). Of the 1.6 Gb assembly, 1.5 Gb could be aligned to the pseudomolecule sequence of asparagus, without any structural rearrangements (Figure 2 and Table 2). Since we determined haplotype-phased genome sequences for the male and female lines of *A. kiusianus*, it was possible to compare the sequence and structure of the *M* locus between the Y-specific region of the male line and the corresponding region of the female line (Figure 3). The result suggested dynamic genome rearrangements between the male and female lines, similar to that reported in jojoba^9^, which might lead to presence/absence variation of the male-specific gene *MSE1/AoMYB35/AspTDF1* between the male and female lines^3–5^.

The genome of *A. kiusianus* harbors valuable genes that could be used for the breeding of elite garden asparagus cultivars. Because of cross-compatibility between the two species^2,10^, important genetic loci such as those imparting resistance to stem blight^2^, which causes considerable production losses, could be transferred from *A. kiusianus* into garden asparagus. However, while DNA markers linked to the genes would facilitate the selection of disease-resistant lines in breeding programs, the genetic loci responsible for disease resistance have not been reported so far. The chromosome-level genome sequence of *A. kiusianus* presented in this study could serve as a reference for genetic mapping and the identification of resistance genes, as well as for transcriptome analysis and the determination of gene functions and mechanisms underlying the resistance and susceptible phenotypes^6,7^.

Although the plant genomics era started with the whole-genome sequencing of *Arabidopsis thaliana*^12^, an undomesticated species, the advanced approaches of plant genomics have been applied more frequently to agronomically important crops rather than to wild plant species^11^. Wild plants have the potential to accelerate the pace of breeding programs and to further the field of plant science^13,14^. The genome sequence information of *A. kiusianus* generated in this study will help to reveal the genetic mechanisms underlying sexual differentiation in plants and will accelerate disease-resistance breeding in asparagus.

## Data availability

Sequence reads generated in this study are available from the Sequence Read Archive (DRA) of the DNA Data Bank of Japan (DDBJ) under accession number: DRA012987. The DDBJ accession numbers of the assembled genome sequences are BQKN01000001-BQKN01111443 (AKIK1p1), BQKO01000001-BQKO01111443 (AKIK1p2), BQKP01000001-BQKP01125241 (AKIK2p1), and BQKQ01000001-BQKQ01125241 (AKIK2p2). The genome sequence information generated in this study is available at Plant GARDEN (https://plantgarden.jp).

## Supplementary data

**Supplementary Table S1** Software tools used for genome assembly and gene prediction.

**Supplementary Table S2** Statistics of the primary genome sequence assemblies of the male and female lines of *Asparagus kiusianus*.

## Acknowledgements

We thank T. Ikeuchi and T. Nakamura (Kagawa Prefectural Agricultural Experiment Station) for providing the plant material, and A. Watanabe (Kazusa DNA Research Institute) for technical assistance.

